# A large effect fitness trade-off across environments is explained by a single mutation affecting cold acclimation

**DOI:** 10.1101/2023.09.11.557195

**Authors:** Gwonjin Lee, Brian J. Sanderson, Thomas J. Ellis, Brian P. Dilkes, John K. McKay, Jon Ågren, Christopher G. Oakley

## Abstract

Identifying the genetic basis of local adaptation and fitness trade-offs across environments is a central goal of evolutionary biology. Cold acclimation is an adaptive plastic response for surviving seasonal freezing, and costs of acclimation may be a general mechanism for fitness trade-offs across environments in temperate zone species. Starting with locally adapted ecotypes of *Arabidopsis thaliana* from Italy and Sweden, we examined the fitness consequences of a naturally occurring functional polymorphism in *CBF2*. This gene encodes a transcription factor that is a major regulator of cold-acclimated freezing tolerance, and resides within a locus responsible for a genetic trade-off for long-term mean fitness. We estimated the consequences of alternate genotypes of *CBF2* on 5-year mean fitness and fitness components at the native field sites by comparing near isogenic lines with alternate genotypes of *CBF2* to their genetic background ecotypes. The effects of *CBF2* were validated at the nucleotide level using gene edited lines in the native genetic backgrounds grown in simulated parental environments. The foreign *CBF2* genotype in the local genetic background reduced long-term mean fitness in Sweden by more than 10%, primarily via effects on survival. In Italy, fitness was reduced by more than 20%, primarily via effects on fecundity. At both sites, the effects were temporally variable and much stronger in some years. The gene edited lines confirmed that *CBF2* encodes the causal variant underlying this genetic trade-off. Additionally, we demonstrated a substantial fitness cost of cold acclimation, which has broad implications for potential maladaptive responses to climate change.

## Introduction

Adaptive differentiation among natural populations is driven by divergent selection. This may lead to local adaptation, where the local ecotype outperforms foreign ecotypes (1, 2). Reciprocal local adaptation in transplant experiments is direct evidence of fitness trade-offs across environments, i.e., that adaptation to one environment reduces fitness relative to local ecotypes in other environments (2). Such fitness trade-offs have long been thought to be important drivers of biological diversification across scales (3–5). Despite numerous studies demonstrating local adaptation (2, 6–8) the genetic and physiological mechanisms underlying local adaptation and fitness trade-offs across environments remain poorly understood (9–12), as does the mechanistic basis of adaptation more generally (13–15). The identification of causal variants for adaptation bears on long-standing questions with important consequences for the rate and predictability of adaptive differentiation, including the role of large effect alleles in adaptation (16–19), and the contribution of individual loci to fitness trade-offs across environments (20, 21).

Despite a large literature on the genetic basis of adaptation, understanding of the full causal chain connecting naturally occurring sequence polymorphism to traits (molecular or organismal), and ultimately to fitness in the natural environments in which the organisms have evolved remains an important and elusive goal. Studies have mapped genetic loci for local adaptation (22–29) but rarely identify and functionally validate causal polymorphisms. The genetic basis of natural variation in traits that are ecologically important under some contexts has been discovered and functionally validated (30–33). However, pervasive genotype by environment interactions for both traits and fitness (21, 31, 34) means that interpreting these results in the context of local adaptation and fitness trade-offs requires testing the natural alleles in the genetic backgrounds in which they occur, and in the native environments in which the organism evolved. Indeed, recent papers on the genetics of adaptation highlight the need to follow up mapping studies with functional validation of candidate genes (15), and the need to explicitly test for fitness effects (19). Answering these calls requires integration of field study of adaptation with experiments in realistic conditions on natural ecotypes with experimentally manipulated alleles. This is currently practical in only a few study systems.

One potential mechanism for fitness trade-offs across environments in broadly distributed temperate zone species is cold-acclimated freezing tolerance. Cold acclimation is a plastic response to low temperatures, and increases survival in climates with freezing winter temperatures in both animals and plants (35–39). Cold acclimation involves dramatic changes in transcription, protein translation, and metabolism in response to cool temperatures in animals (40–44) as well as plants (37, 38, 45). Cold acclimation is thought to be energetically costly (46), and could lead to broad-scale trade-offs when cool temperatures that induce this response are more geographically widespread than is the occurrence of freezing temperatures. Producing the wrong phenotype for a given environment is one possible cost of plasticity, though additional costs of plasticity such as maintaining the machinery to sense and respond to variation have also been proposed (47). In animals, a cost of cold acclimation has been demonstrated in *Drosophila* (48–50). Cold acclimation may also lead to maladaptive responses if cues earlier in the season become poorer predictors of the severity of subsequent winter because of climate change and climate variability.

In winter annuals and other plants, cold acclimation can increase freezing tolerance through the accumulation of soluble sugars such as raffinose (51–53) and other compounds that help decrease the freezing point of the cell, resist desiccation, and stabilize and protect cell membranes (37, 38, 45, 54). In *Arabidopsis thaliana*, C-repeat/DRE binding factor 2 (CBF2) is a member of the Apetala2 family of transcription factors and is a necessary regulator of cold acclimation in laboratory studies (45, 55). Natural variation in *CBF2* contributes to differences in freezing tolerance across *A. thaliana* ecotypes (56–59). Cold responsiveness of CBF orthologs appears to be broadly conserved in plants (37, 38), and contributes to freezing tolerance in diverse species such as canola (60), wheat (61), and poplar (62), so it seems likely that this specific mechanism of regulating cold acclimation is of general importance. Cold acclimation is common in temperate zone plants, and more information on the costs of cold acclimation can contribute to our understanding of genetic trade-offs across environments and the potential negative fitness consequences of climate change.

Testing for costs of cold acclimation in plants is challenging and results are inconclusive thus far. Experimental tests on tree ecotypes from different geographic origins under different experimental conditions found that it is difficult to tease apart acclimation responses from phenological adaptations that are also cued by temperature and photoperiod (63). Studies in *A. thaliana* have shown that overexpression of *CBF2* leads to a stunted phenotype and reduced fitness (64, 65), providing indirect evidence of a cost of *CBF2* mediated cold acclimation. Clinal and correlational associations also suggest a potential cost of cold acclimation (66–69). While the main benefit of cold acclimation in freezing environments is increased survival, it is unknown whether the costs of acclimation will be manifest in reduced survival, fecundity, or both. To our knowledge there has only been one direct test of the costs of cold acclimation in plants, which reported no cost for short term acclimation of *Arabidopsis thaliana* in the lab (65). However, the costs of cold acclimation may be expressed only under ecologically relevant conditions (c.f., 48, 49).

In our study system of locally adapted ecotypes of *Arabidopsis thaliana* from Sweden and Italy, cold acclimation likely plays a major role in fitness trade-offs across environments (29, 57, 59, 70). A correlation between minimum winter temperature and relative survival of the Italian ecotype in Sweden indicated that sub-freezing temperature is a primary selective agent in Sweden (57, 70). At the field site in Italy, autumn and winter temperatures are cool but typically non-freezing (57, 71), so the cost of acclimation will be expressed without the benefit. Furthermore, major freezing tolerance quantitative trait loci (QTL) co-localize with fitness trade-off QTL detected over multiple years at the native field sites (29, 57). This strongly suggests that cold acclimation mediated by these freezing tolerance loci are a mechanistic basis of genetic trade-offs. The causal variant for the largest effect freezing tolerance QTL is a loss of function mutation in the Italian allele of *CBF2* (58), which can explain 1/3 of the difference in freezing tolerance between the Swedish (SW) and Italian (IT) ecotypes (59).

The final step in conclusively demonstrating a role of *CBF2* in a fitness trade-off is to link the effects of this naturally occurring *CBF2* polymorphism to lifetime fitness in both environments. We predicted that the functional Swedish genotype of *CBF2* will be adaptive in Sweden, but that cold acclimation induced by a functional *CBF2* in Italy will incur a fitness cost. As EU regulations prohibit planting GMO lines in the field, we took a two-step approach. We used Near Isogenic Lines (NILs) in field experiments, and gene edited lines with manipulated alleles of *CBF2* in the native genetic backgrounds in growth chambers programmed to mimic temperature and photoperiod at the native sites. We addressed the following questions: 1) What are the effects of introgression segments containing alternate alleles of *CBF2* on estimates of long-term mean fitness when grown at the native sites? 2) What are the effects of this single gene on fitness and fitness trade-offs across environments? 3) Are the benefits and costs of *CBF2* in alternate environments mediated by viability selection, fecundity selection, or both?

## Results

### Reciprocal transplant experiments at the Swedish and Italian field sites – Overall fitness

There was a strong and statistically significant fitness tradeoff across environments between the ecotypes for estimates of mean lifetime fitness (total fruit production including zeros for plants that died without having produced any fruits) over five years (Fig. 1; Table S1). This result is consistent with previous findings in this system (29, 70), and is an important prerequisite for investigating the contribution of any polymorphism to local adaptation and fitness trade-offs across environments. In Sweden, five-year mean fitness of the Italian ecotype (IT) was 65% less than that of the local Swedish (SW) ecotype (Fig. 1A), and in Italy five-year mean fitness of the SW ecotype was 82% less than that of the local IT ecotype (Fig. 1B). Thus, there is a pattern of strong overall selection against the foreign ecotype at both sites, though the strength of selection was somewhat stronger in Italy than in Sweden.

**Figure 1.**
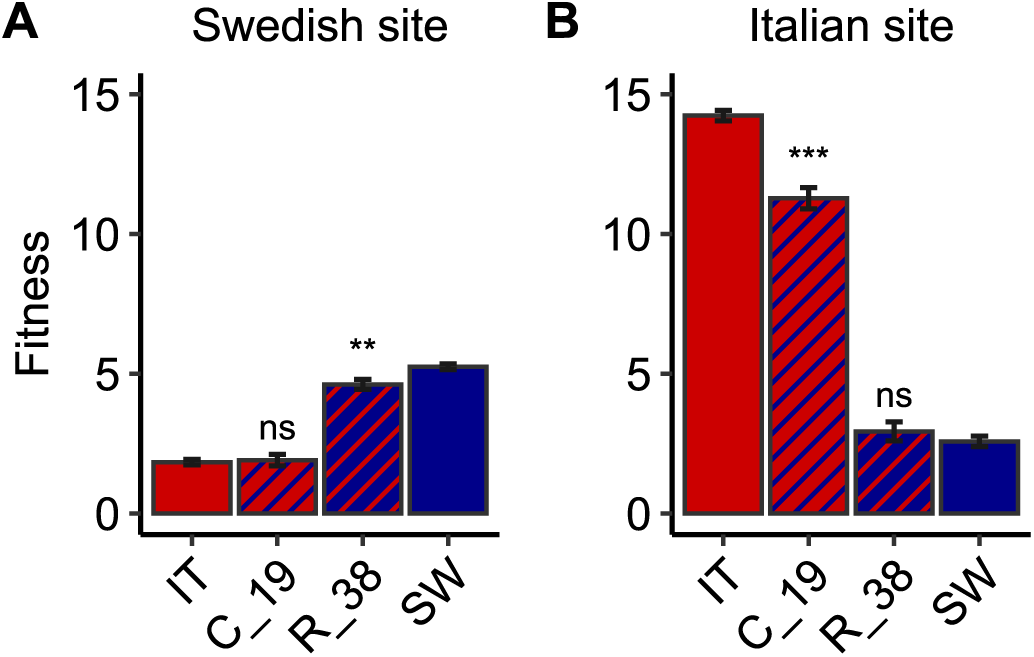
Least squares mean estimates of fitness (number of fruits per seedling planted) of the SW and IT ecotypes and NILs over five individual one-year experiments at the Swedish (A) and Italian (B) field sites. Error bars are 1 SE. The predominant color indicates the genetic background (SW ecotype = blue; IT ecotype = red). Hatching indicates the genotype of the introgression segment containing *CBF2* in the NILs. Asterisks represent statistically significant contrasts between a NIL and its genetic background (***, *P* < 0.001; **, *P* < 0.01; ns, not significant).

To quantify the effect of *CBF2* at each site, we used Near Isogenic Lines (NILs) with introgression segments containing alternate genotypes of *CBF2* in each genetic background (Fig. S1) and compared their 5-year mean fitness to that of their genetic background ecotype. There was a strong and statistically significant signal of a genetic trade-off across environments for the foreign genotypes of *CBF2* in the native genetic backgrounds (Fig. 1; Table S2). At the Swedish site, the NIL containing the IT *cbf2* loss-of-function (LOF) genotype in the SW genetic background had 12% lower fitness than the SW ecotype. At the Italian site, the NIL containing the functional SW *CBF2* genotype in the IT genetic background had 21% lower fitness than the IT ecotype. NILs with the local *CBF2* genotype in the foreign genetic background had a 4% and 14% increase in fitness in Sweden and Italy, respectively, though these effects were not statistically significant.

Within individual years there were consistent differences between ecotypes reflecting local adaptation, but the effects of the introgression segments containing *CBF2* were temporally variable at both sites (Fig. 2; Tables S1 & S2). Focusing on the foreign *CBF2* genotype in the local genetic background, we found a statistically significant effect in two out of five years in Sweden, with reductions in fitness of up to 71%. In Italy, there were significant effects in three years (with a suggestive effect in a fourth year), with reductions in fitness of up to 40% (Table S2). Effect sizes in the remaining non-significant contrasts were mostly, but not always, weak.

**Figure 2.**
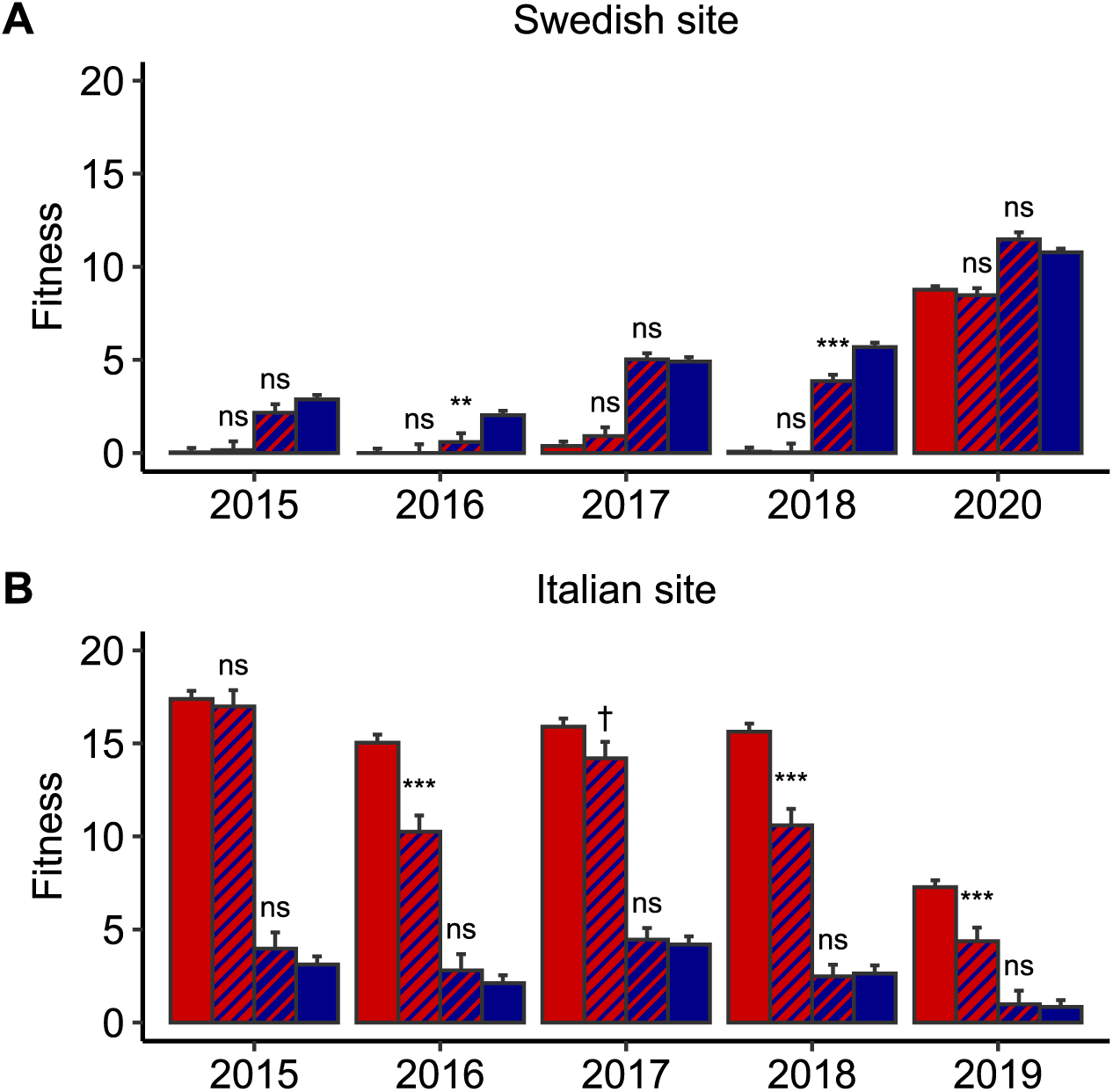
Least squares mean estimates of fitness (number of fruits per seedling planted) of the ecotypes and NILs in each individual year at the two field sites. Error bars are 1 SE. The predominant color indicates the genetic background (SW ecotype = blue; IT ecotype = red). Hatching indicates the genotype of the introgression segment containing *CBF2* in the NILs as shown in Figure 1. Asterisks denote significant contrasts between a NIL and its genetic background (***, *P* < 0.001; **, *P* < 0.01; ^†^, 0.05 < *P* < 0.1; ns, not significant).

### Single gene effects of *CBF2* in simulated Italian and Swedish conditions – Overall fitness

To test the effects of *CBF2* at the single gene level required us to use growth chamber experiments simulating key environmental features at the native sites (Fig. S2). The growth chamber programs (Fig. S3; Supporting text) were constructed based on long-term climate data collected at the native sites. These programs were then tested and refined to recapitulate differences in relative fitness between ecotypes observed at these sites (Fig. S4). The growth chamber experiments included the parental ecotypes, two independent CRISPR induced LOF *cbf2* lines in the SW genetic background, and two transgenic lines containing SW *CBF2* and native promoter in the IT background. We omitted the IT background lines from the Swedish chamber experiment because the very low fitness of IT background lines in the field experiment (Fig. 1) and results of freezing tolerance assays with these lines (59) suggested it would not be possible to detect effects in this background in the Swedish environment.

There was a strong signature of a fitness trade-off across environments between the ecotypes in the growth chambers, and single gene estimates of the effects of *CBF2* corroborate the causality of this gene for the NIL effects estimated in the field. We found very strong and statistically significant selection against the IT ecotype in the Swedish chamber (92%) and the SW ecotype in the Italian chamber (62%; Figs. 3 & S4; Table S3). Moreover, in the Swedish chamber, both independent LOF *cbf2* lines in the SW background had significantly reduced fitness (23% and 22%) compared to the SW ecotype (Fig. 3A; Table S2). In the Italian chamber, both independent transgenic SW *CBF2* lines in the IT background had significantly reduced fitness (12% and 21%) compared to the IT ecotype, and the lines with IT *cbf2* in the SW background had significantly greater fitness (23% and 16%) than the SW ecotype (Fig. 3B; Table S2).

**Figure 3.**
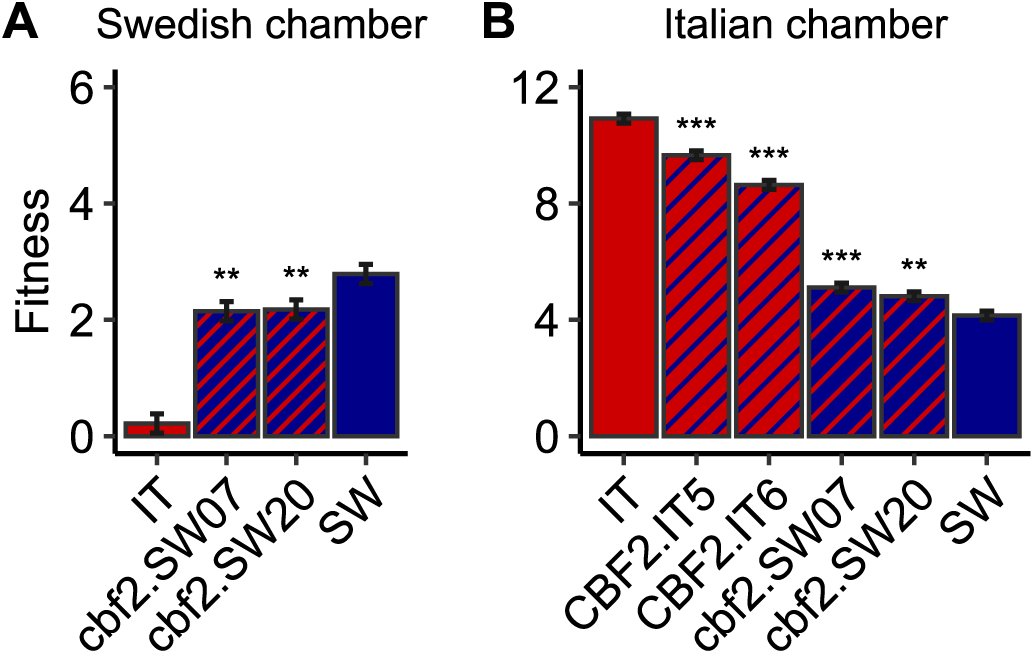
Least squares mean fitness (number of fruits per seedling planted) of the ecotypes and gene edited lines in the “Swedish” (A) and “Italian” (B) growth chambers. Error bars are 1 SE. The predominant color indicates the genetic background (SW ecotype = blue; IT ecotype = red). Hatching indicates the genotype of *CBF2* (capitals = functional copy, lower case = loss of function) in gene edited lines. Asterisks represent statistically significant contrasts between a gene edited line and its genetic background (***, *P* < 0.001; **, *P* < 0.01). Note that the scale of the Y axis of fitness for the Italian growth chamber is double that of the Swedish growth chamber.

### Reciprocal transplants at the Swedish and Italian field sites – Fitness components

To determine if the costs (Italy) and benefits (Sweden) of functional CBF2 are mediated through viability selection, fecundity selection, or both, we compared survival and fecundity of the lines in the field experiment. The ecotypes were significantly locally adapted for both fitness components in Italy, and for survival in Sweden (Fig. 4; Table S1). Note that because of very low survival of the Italian ecotype at the Swedish site, it was only possible to compare the fecundities of the two ecotypes in one of five years (Fig. S5; Table S2). The IT ecotype had 67% lower survival than SW in Sweden, and SW had 33% lower survival and 73% lower fecundity than IT in Italy. The contribution of *CBF2* to fitness trade-offs was most pronounced for survival in Sweden and fecundity in Italy (Figs. 1 & 4; Table S2). For survival in Sweden, the NIL with IT *cbf2* in the SW background had 13% reduced survival compared to SW, and SW *CBF2* in IT had 15% increased survival compared to IT, though the latter difference was only suggestive (Fig. 4A; Table S2).

**Figure 4.**
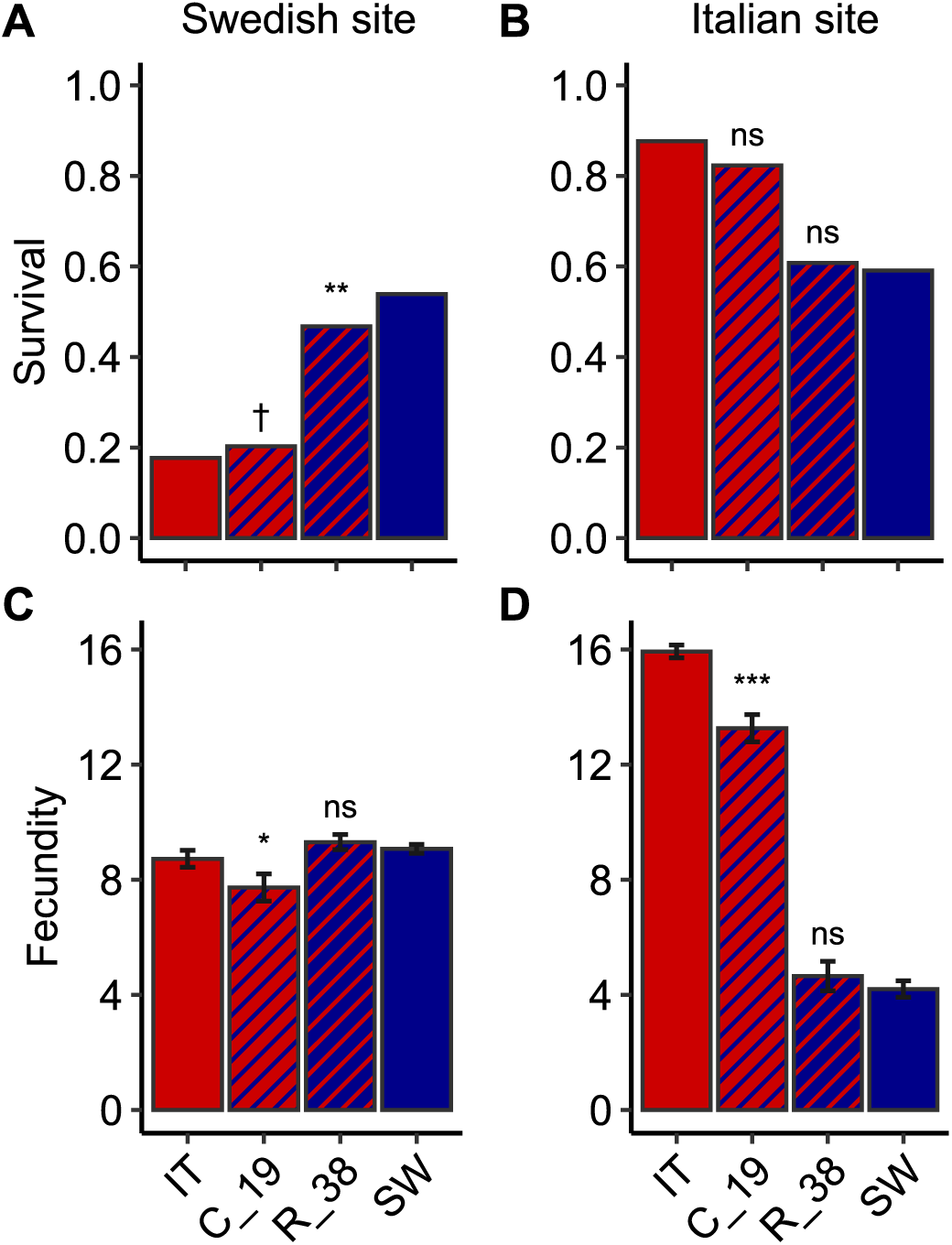
Fitness components of the SW and IT ecotypes and NILs over five individual one-year experiments at the Swedish and Italian field sites. Mean proportion survival (averaged over years) at the Swedish (A) and Italian (B) field sites. Least squares mean fecundity at the same sites (C and D). Colors and hatching as in other figures. Error bars for fecundity are 1 SE, no bars are given for survival because it was analyzed with a binomial error distribution. Asterisks represent statistically significant contrasts between a NIL and its genetic background (***, *P* < 0.001; **, *P* < 0.01; *, *P* < 0.05; ^†^, 0.05 < *P* <0.1; ns, not significant).

Effects of IT *cbf2* in SW on fecundity in Sweden were weak and non-significant. Functional *CBF2* in IT decreased fecundity by 11% (Fig. 4C; Table S2), but both the sign and magnitude of this effect must be interpreted cautiously because of extremely high mortality of IT in all years except 2020. In Italy, effects of *CBF2* on survival were weak and not statistically significant in both backgrounds (Fig. 4B; Table S2). For fecundity in Italy, functional *CBF2* in IT reduced fecundity by 17% compared to IT and non-functional *cbf2* in SW increased fecundity by 11%, though only the former contrast was statistically significant (Fig. 4D; Table S2).

Within individual years there were consistent differences between ecotypes for fitness components reflecting local adaptation except for fecundity in Sweden (Fig. S5, Table S1), where in many years too few IT individuals survived to make comparisons of fruit production to other genotypes meaningful. For survival in Sweden and both fitness components in Italy, the effects of *CBF2* varied across years. We focus on the effects of the foreign *CBF2* genotype in the local genetic background compared to the local ecotype. For survival, there were statistically significant reductions (up to 45%) in Sweden in two years with a suggestive effect in a third year; in Italy survival was significantly reduced in one year (Figure S5; Table S2). For fecundity in Italy, there were significant decreases (up to 31%) in two years (Figure S5; Table S2).

### Single gene effects of *CBF2* in simulated conditions – Fitness components

Patterns of relative survival and fecundity of the ecotypes, and the effects of *CBF2* on those fitness components in the growth chamber experiments were consistent with the field results. In the Swedish environment, effects were recorded primarily on survival, whereas in the Italian environment, effects were recorded primarily on fecundity (Figs. 3 & S6; Table S3). In the Swedish chamber, the survival of the IT ecotype was 91% lower than that of the SW ecotype, whereas in the Italian chamber, SW had 62% lower fecundity than IT. In the Swedish chamber, both independent LOF *cbf2* lines in the SW background had strongly and significantly reduced survival compared to the SW ecotype (Fig. S6A; Table S2). In the Italian chamber, both independent transgenic SW *CBF2* lines in the IT background had significantly reduced fecundity compared to the IT ecotype, and the lines with IT *cbf2* in a SW background had significantly greater fecundity than the SW ecotype (Fig. S6D; Table S2).

## Discussion

Identifying the genetic basis of local adaptation and fitness trade-offs across environments is a central, but often unrealized goal in evolutionary biology. We demonstrated that reciprocal introgressions of a genomic segment containing *CBF2* strongly affected the long-term mean fitness of two locally adapted ecotypes of *A. thaliana* when grown in a 5-year reciprocal transplant experiment conducted at the native sites. We then functionally validated that *CBF2* is the causal locus for this variation using replicated single gene edited lines in the native genetic backgrounds grown in simulated parental environments. Taken together, the results provide a unique confirmation of a causal polymorphism underlying local adaptation, and demonstrate a substantial cost of cold acclimation with broad implications for potential climate-change induced maladaptation of temperate zone plants.

At both sites, we found strong effects of introgression segments containing *CBF2* on fitness, but these effects were temporally variable and depended on genetic background. These results confirm the strong and temporally variable effect of the genomic region around *CBF2* first reported from QTL mapping over multiple years at the same sites (26, 29). Despite the effort of replicating the present field experiment for five years, correlations between effect sizes and climate variables with five data points per site would be tenuous. However, the two years with significant effects of *CBF2* on fitness in Sweden had the coldest winter minimum soil temperatures, consistent with a role of *CBF2* in freezing tolerance (57, 59). In Italy, where freezing soil temperatures are extremely rare, the two years without significant effects of *CBF2* on fitness had warmer mean soil temperatures during the coldest period. This suggests greater costs of inducing cold acclimation during cooler winters in Italy. There are at least two possible explanations for the tendency for *CBF2* effects to be statistically significant only in the local genetic background. It may simply be that the very poor fitness of the foreign ecotype makes it more difficult to detect a significant effect. Alternatively, there may be epistatic effects between CBF2 and downstream target genes, but to test this hypothesis requires additional data.

Together, these results indicate that the effects of a genetic trade-off can be context dependent, and the magnitude and ability to detect an effect at a locus will depend on the genetic background, as well as spatial and temporal environmental variation.

Comparison of effect sizes between the current NIL experiment and previous QTL experiments is difficult, both because the experiments took place mostly in different years, and because the effect sizes represent different contrasts. Here effect sizes represent the effect of a single segment tested against an isogenic ecotypic background, but effect sizes in the QTL study represent the effect of different alleles at a SNP averaged over heterogenous RIL backgrounds and tested as a deviation from the mean fitness of all RILs. That said, the effects of this genomic region in the NIL experiment seem to be greater than those in the QTL experiment for long-term mean fitness in Italy, and for maximum annual effects at both sites. This suggests that estimates of effect sizes previously reported for this QTL are not inflated as might be expected by the “Beavis effect” (72), and our results are among a growing number of examples of large effect loci involved in adaptation and adaptive traits (18, 19, 73, 74). Transcription factors may be particularly likely candidates for large effect loci involved in broad scale adaptation because they have downstream effects on the expression level of many other genes (75, 76). While the field experiments in the native environments replicated over multiple years represent the most natural test of the fitness consequences of alternate alleles of *CBF2*, an important caveat about these results is that the introgression segments in the NILs may well include polymorphisms in additional genes affecting fitness.

Raising ecotypes and gene edited lines in growth chamber programs that mimic temperature and photoperiod changes at the two sites was therefore a critical next step in functional validation of the *CBF2* polymorphism. Estimates of the relative fitness of the two ecotypes in the growth chamber experiments were qualitatively similar to those in the field experiments, demonstrating that the growth chambers replicated some essential features of the field environments (Figs. 1, 3, & S4). Moreover, the effects of single gene edits of *CBF2* in the local genetic background on both cumulative fitness and fitness components (Figs. 3 & S6) were qualitatively similar to mean effects observed in the 5-year field experiment using NILs (Figs. 1 & 4), and well within the range of effect sizes observed in individual years in the field (Fig. 2 & S5). The results conclusively demonstrate a causal role for *CBF2* in the fitness trade-off observed across the growth chamber environments, and are consistent with a similar role for the trade-off across the two native sites.

Elucidating the genetic basis of local adaptation is challenging. Few systems allow for direct functional validation of the fitness consequences of naturally occurring sequence polymorphisms in the native genetic backgrounds grown in their native environments. Studies mapping QTL for local adaptation in field reciprocal transplants (reviewed in 21, 28, 29) identify naturally occurring polymorphisms in regions of the genome associated with fitness, but these QTL often contain many genes that differ between the founder parents. Genome-wide association mapping (GWAS) of the genetic basis of regional adaptation can provide better, but often still incomplete, resolution of candidate polymorphisms (77–79). Both approaches test genetic hypotheses, but the molecular identities of the causal genes need to be tested by functional validation (11, 13, 15). Such validation is limited to systems in which genetic manipulation is feasible, or where one can indirectly test the effect of a gene from the focal species in a model organism (80, 81). Functional genetic studies on putatively adaptive traits provide a clear link between causal variant and phenotype (30–33). Other work has evaluated the fitness effects of mutations in field experiments (75, 82). We extended this approach to a reciprocal transplant in which the alleles affecting a fitness trade-off were evaluated in the native genetic backgrounds and their environments. The relationships between genotype, phenotype, and fitness depend on the context of the environment and genetic background (19, 21, 31, 34). Our approach puts the genetic effects in such context, though legal constraints necessitated evaluating the single gene edits in simulated rather than actual parental environments. The concordance between field and growth chamber results demonstrate that we captured some essential differences observed in nature.

The mechanistic basis of the fitness effects of this functional polymorphism in the Swedish environment stems from the major role of the CBF2 transcription factor in the regulation of cold acclimation. We hypothesize that investment in acclimation in response to cool, but non-freezing temperatures, results in the fitness cost in the Italian environment. This polymorphism in *CBF2* has previously been shown to explain about 1/3 of the difference between the SW and IT ecotypes in cold-acclimated freezing tolerance in the lab (57, 59), and the effects of natural variation in *CBF2* on freezing tolerance has been demonstrated in other ecotypes as well (56). In our system, a short list of candidate genes that are partially regulated by CBF2 in response to short-term cold acclimation have been identified using RNAseq (59). These genes have likely roles in sugar biosynthesis, desiccation resistance, and membrane stabilization. The strongest of these candidates was *Galactinol synthase 3* (*GolS3*), which is an important enzyme in the synthesis of galactinol and raffinose. Accumulation of raffinose and other soluble sugars has previously been correlated with increased freezing tolerance (51, 52). In *A. thaliana*, accessions from higher latitudes have greater cold acclimated freezing tolerance (67, 83). Accessions from colder climates also accumulate more raffinose in response to cold, and this is negatively correlated with vegetative growth rates (53). Additional work is needed to characterize the role of the *CBF2* polymorphism on transcriptional and metabolic responses across the cold periods in both environments to better characterize the contribution of raffinose and other compounds to fitness trade-offs across environments.

Regardless of the specific mechanisms, our fitness results in the Italian environment provide direct evidence for a cost of cold acclimation, in agreement with several prior lines of indirect evidence. We hypothesize that such a cost is borne even in the Swedish environment, but that in most years the fitness benefits of freezing tolerance outweigh the costs of cold acclimation. This could have profound implications for maladaptive responses in many temperate zone plants with climate change. A decoupling between cold acclimation cues and the severity of subsequent winter could lead to reductions in fitness if autumn temperatures are insufficient to acclimate for an anomalously severe winter, or if cool autumn temperatures that induce cold acclimation are followed by a milder winter. Direct evidence for a cost of cold acclimation in plants has until now proven difficult. This is likely both because manipulating the acclimation environment (and/or using a broad geographic sample of ecotypes) also affects phenological patterns in species where cold serves to cue phenological transitions (63), and because costs of acclimation are context-dependent. Accurate quantification of the potential costs of cold acclimation will likely require manipulating major regulators of cold acclimation as we have done here. Orthologs of CBF genes are cold responsive in many plant lineages (37). Our approach could be applied more broadly as transformation becomes feasible in more species. In the meantime, we suggest that predictions about organismal resilience in the face of climate change should carefully consider potential costs of cold acclimation.

## Materials and Methods

### Study system

*Arabidopsis thaliana* is an annual, predominantly selfing plant native to habitats in Eurasia and Africa (84, 85). Our source ecotypes are from locally adapted populations from Rödåsen (hereafter “SW”) in north-central Sweden (62°48′N, 18°12′E) and Castelnuovo di Porto (hereafter “IT”) in central Italy (42°07′N, 12°29′E), near the northern and southern edge of the native Eurasian range (70). At both sites, plants exhibit a winter annual life history and overwinter as vegetative rosettes. At the Italian site (42°07′N, 12°29′E), seeds germinate in autumn, and plants overwinter as rosettes under cool conditions followed by flowering in March and April. At the Swedish site (62°48′N, 18°12′E), seeds germinate in late summer, and plants overwinter as rosettes and flower in May and June. These ecotypes have been used extensively to map the genetic basis of local adaptation (26, 27, 29, 86, 87) and ecologically important traits (27, 57, 88, 89), as well as to study physiological responses that are infeasible to map (90, 91).

### Development of genetic resources

To estimate lifetime fitness effects of *CBF2* in both native environments, we used a combination of Near Isogenic Lines (NILs) and gene edited lines. Details of NIL construction can be found in (Fig. S1) and (59). We used two NILs containing alternate introgression segments surrounding *CBF2* in each genetic background. The SW background NIL contains a segment with the IT loss-of-function (LOF) *cbf2* genotype. The IT background NIL contains a segment with the functional SW *CBF2* genotype (Fig. S1). These segments contain many genes in addition to *CBF2* and cannot provide single gene resolution, but were used to quantify fitness effects of this genomic region in field experiments at the native sites where European Union regulations prohibit the planting of GMO organisms.

To obtain single gene resolution of the fitness effects of *CBF2* we used gene edited lines and the parental ecotypes in growth chamber experiments. To mimic the IT *cbf2* LOF ecotype in the SW genetic background we used two independent *cbf2* LOF lines (produced using CRISPR/Cas9). These lines, as well as NILs, have previously been shown to explain over 1/3 of ecotypic differences in cold acclimated freezing tolerance (59). For IT background lines with a functional copy of *CBF2*, we used two independent transgenic lines with SW *CBF2* and native promoter inserted into the IT background. These lines were previously used in electrolyte leakage assays of freezing tolerance (58).

### Reciprocal transplant experiments at the Swedish and Italian field sites

To estimate the effects of introgression segments containing alternate genotypes of *CBF2* in both genetic backgrounds, we quantified fitness and fitness components in field reciprocal transplant experiments. We transplanted seedlings of the two ecotypes and two NILs (R_38 and C_19; Fig. S1) at both the native Swedish and Italian field sites in each of five years. Seed germination, transplanting, and field planting protocols closely follow previous field studies of these ecotypes (26, 29). In brief, in each individual year surface sterilized seeds were sown on agar in Petri dishes, and cold-stratified in the dark at 4°C for 1 w to break seed dormancy and synchronize germination. The Petri dishes were then transferred to a growth room set to 22°C, 16-h day (16L:8D) with photosynthetically active radiation (PAR) of 150 µmol photons m^-2^ s^-1^ for eight to ten days for germination. Seedlings were transplanted into a randomized design in plug trays with individual cells of 20 x 20 x 40 mm filled with an equal mixture of local sand, gravel, and unfertilized peat in Sweden and with local soil in Italy. In each of five years, trays with seedlings were then set into the ground in early September at the Swedish site, and in early November at the Italian site. Sample sizes per genotype per year were variable across years but ranged from 400-600 seedlings each for IT (mean=438) and SW (mean=440), 100-150 seedlings for C_19 (mean=109), and 100-200 seedlings for R_38 (mean=150). For each individual plant we estimated cumulative fitness as the total number of fruits produced per seedling planted, which incorporates both survival and fecundity (fruit number per surviving plant). Logistical constraints prevented us from quantifying seed number per fruit to obtain a more complete estimate of cumulative fitness, but previous work in this system has shown that estimating fitness as fruits per seedling and seeds per seedling yield qualitatively similar patterns (87). Data from the present experiment is distinct from a recent reciprocal transplant at the same sites (29), though parts of both experiments were conducted in 2016 and 2017.

Within each site, we examined the effects of genotype, year and the genotype x year interaction on cumulative fitness, survival, and fecundity using ANOVA. Both genotype (two ecotypes and two NILs) and year were treated as fixed effects. Survival was analyzed with a binomial error distribution. Inspection of residuals for fecundity indicated that a normal error distribution was suitable. For cumulative fitness there were some departures from normality and equal variances, however overall results using a normal error distribution were qualitatively similar to non-parametric models and models with alternative error distributions (including zero inflated Poisson and zero inflated negative binomial), so we present the normal model results for simplicity. For any model with a significant effect of genotype or genotype x year, we used *a priori* linear contrasts to compare the IT and SW ecotypes. We also tested for the effect of the *CBF2* region on differential adaptation by contrasting NILs to their genetic background ecotypes.

### Single gene effects of CBF2 in simulated Italian and Swedish conditions

To quantify the fitness effects of the single *CBF2* polymorphism, we developed growth chamber programs to simulate seasonal changes in temperature and photoperiod at the Swedish and Italian sites. For details of field climate data see Fig. S2 and (71). Daily mean high and low temperature values across years were used to guide construction and optimization of the growth chamber programs (Fig. S3, Supporting text). Although it is impossible to completely mimic field conditions in a growth chamber, the general concordance in ecotypic differences in relative fitness between field and growth chamber experiments (Fig. S4) gives us confidence that we were able to recreate some of the essential features driving local adaptation in this system.

The optimized growth chamber programs were used to estimate fitness, survival, and fecundity for the IT and SW ecotypes, and gene edited lines with manipulated *CBF2* alleles in the native genetic backgrounds. In both environments, we included two independent *cbf2* loss-of-function lines in a SW background. In the Italian environment, we included two additional transgenic lines containing the SW *CBF2* allele and native promoter in an IT background. The transgenic lines were omitted from the Swedish growth chamber assay because preliminary trials indicated that fitness of IT was so low as to preclude detection of an effect in an IT background line.

Methods for seed sterilization and germination closely follow those of previous freezing tolerance assays in this system (57, 59). In brief, surface-sterilized seeds were sown on agar in Petri dishes. The seeds were cold-stratified in the dark at 4°C for 5 d to synchronize germination. The Petri dishes were then transferred to a growth chamber set to 22°C, 16-h day (16L:8D) with PAR of 125 µmol photons m^-2^ s^-1^ for germination for 12 d. Seedlings were then transplanted into a randomized design in plug trays with individual cells of 20 x 20 x 40 mm filled with propagation mix soil. The trays were then transferred to a growth chamber, LTCB-19 (BioChambers, Winnipeg, MB, Canada) programmed as described above for either the “Italian” or “Swedish” environment (Fig. S3). The entire experiment consisted of 8 trays, arranged side to side on two shelves within the growth chamber. On a given shelf, two border rows around the perimeter of each shelf were planted with “extra” plants (no data collected) to reduce edge effects within the experiment. In the Swedish growth chamber, we added a thin layer of shaved ice onto the plants as soon as chamber temperatures first dropped to freezing conditions (from 4°C to –2°C) to facilitate ice nucleation (58). In each chamber environment, we estimated cumulative fitness for each individual plant as the total number of fruits produced, including zeros for plants that did not survive to reproduce.

Statistical analyses of the growth chamber experiments closely follow that of the field experiments. In the models for the chamber analyses, we also included the effect of tray to account for microspatial variation within the chamber. This was treated as a fixed effect because of the small number of trays. Error distributions for fitness, survival, and fecundity were the same as for the analyses of the field data. In models with a significant effect of genotype, we examined *a priori* contrasts of the IT and SW ecotypes, and contrasted all gene edited lines to their genetic background ecotypes. All statistical analyses for both field and growth chamber experiments were conducted using JMP version 16.

## Acknowledgments

The authors thank dozens of people for assistance with the field and growth chamber experiments including C. Amman, E. Amman, P. Gómez-Zapata, M. I. Jameel, J. Kraft, L. Molina, K. Palacio-López, D. Schemske, F. Spada, M. Vass, L. Vikström, and G. Zacchello. We also thank M. Thomashow for providing seeds of transgenic and CRISPR lines, P. Falzini and Y. Jonsson for permission to conduct experiments on their land, and the Orto Botanico di Roma for allowing us to use their greenhouse facilities. This study was financially supported by grants from the National Science Foundation (DEB-1743273 to CGO and JKM, & IOS-2246545 to CGO and BPD) and the Swedish Research Council (2016-05435, and 2020-04434 to JÅ).

## Supporting Information for

### This PDF file includes

Supporting text

Figures S1 to S6

Tables S1 to S3

SI References

### Supporting Information Text

**Construction and optimization of growth chamber programs to simulate the Swedish and Italian environments.** Growth chamber programs were initially created based on daily mean high and low temperature profiles at the two sites (Fig. S2), and then optimized to satisfy logistical constraints and to recreate ecotypic differences in relative fitness observed at the native field sites. We shortened the overall length of the programs to minimize strain on growth chambers running low temperatures for extended periods. For the Swedish chamber program (Fig. S3A), we shortened the total freezing period and added two cold spikes representative of temperatures in a cold year. For the Italian chamber program (Fig. S3B), this shortening was applied approximately evenly across the program. However, we ensured at least eight weeks of daily high temperatures below 10°C to satisfy the vernalization requirement of the SW ecotype as would occur at the Italian field site. Selection against the foreign ecotype in Sweden has been shown to be positively correlated with winter minimum temperature (1, 2). Photoperiod in the chamber programs was set to reflect seasonal changes at the two sites as described in (3). It is not practical to mimic fluctuation in light intensities observed in the field, so we used a maximum value of photosynthetically active radiation (PAR: µmol m⁻² s⁻¹) of 300 during high temperatures of 19°C or greater, and reduced PAR by approximately 50 units for every reduction in high temperature of ∼3.5°C, similar to our previous protocol (3). During freezing periods in the Swedish chamber, PAR was reduced to zero to minimize circadian effects on freezing tolerance (4, 5) and because daylength is short with low light intensities during this period in the field. Finally, we applied a sequential dry down by gradually reducing the amount of water given at the end of each chamber program to mimic the decreasing soil water potentials observed in the field (6).

Growth chamber programs (Fig. S3) were then fine-tuned to reproduce, as closely as possible, ecotypic differences in relative fitness in field experiments, and ensure that absolute fitness values of the parental ecotypes were within the range observed in the field across all years (1,7). Optimization was done by qualitatively assigning greater weight to temperature profiles of individual years with stronger selection against the foreign ecotype.

**Fig. S1.**
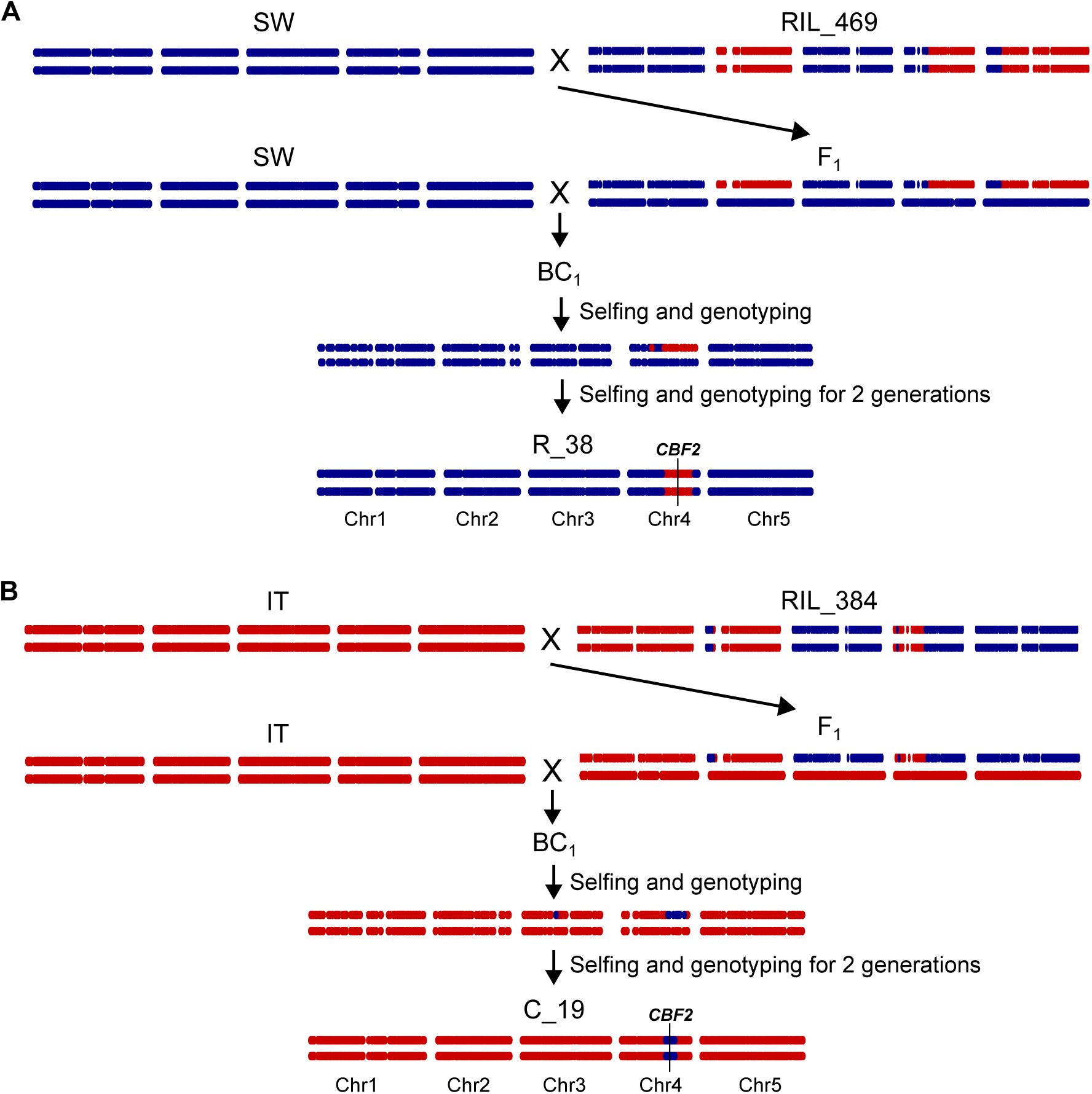
Construction of the SW background NIL (R_38) containing an IT fragment including the *cbf2* LOF mutation (A) and the IT background NIL (C_19) containing a SW fragment including functional *CBF2* (B). Blue points indicate SW marker genotypes and red points are IT marker genotypes (approximately 2000 SNPs). Position of *CBF2* is marked with a black vertical line. NIL construction was initiated by crossing individual recombinant inbred lines (RILs) to the parental ecotypes, and genotyping to confirm formation of an F_1_. The F_1_ was then backcrossed (BC_1_), and resultant progeny genotyped, to isolate alternate introgression segments surrounding *CBF2* on chromosome 4 in otherwise isogenic parental backgrounds. Repeated selfing and genotyping for several generations produced NILs with a single homozygous introgression segment. Genotyping was performed between generations using a combination of 2b-restriction site associated DNA (2b-RAD) sequencing and cleaved amplified polymorphic sequence (CAPS) markers. The SW background NIL (R_38) contains an approximately 8.0-Mb introgression segment. The IT background NIL (C_19) contains an approximately 2.5-Mb introgression segment.

**Figure S2.**
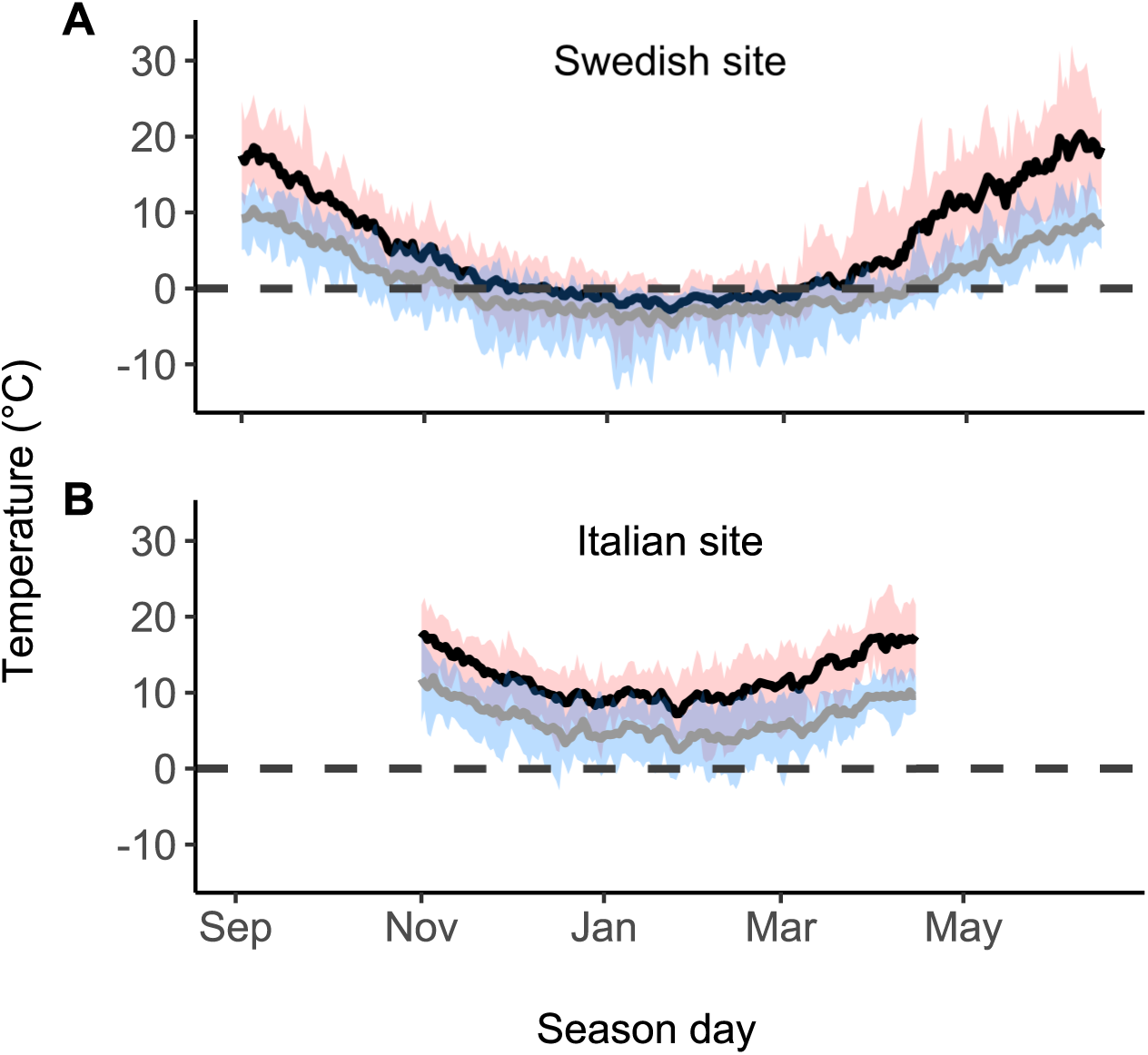
Temperature profiles for the Swedish (A) and Italian (B) field sites from approximately seed emergence to seed production. The solid black and gray lines represent the mean daily maximum and minimum temperatures, respectively across 13 years (2003–2015). The pink and blue shaded regions represent the absolute range of daily maximum and minimum temperatures (averaged over soil and air temperatures) observed across the years. For each day at each site, we averaged over multiple time periods for each sensor, and then averaged over sensors for both air and soil temperature.

**Figure S3.**
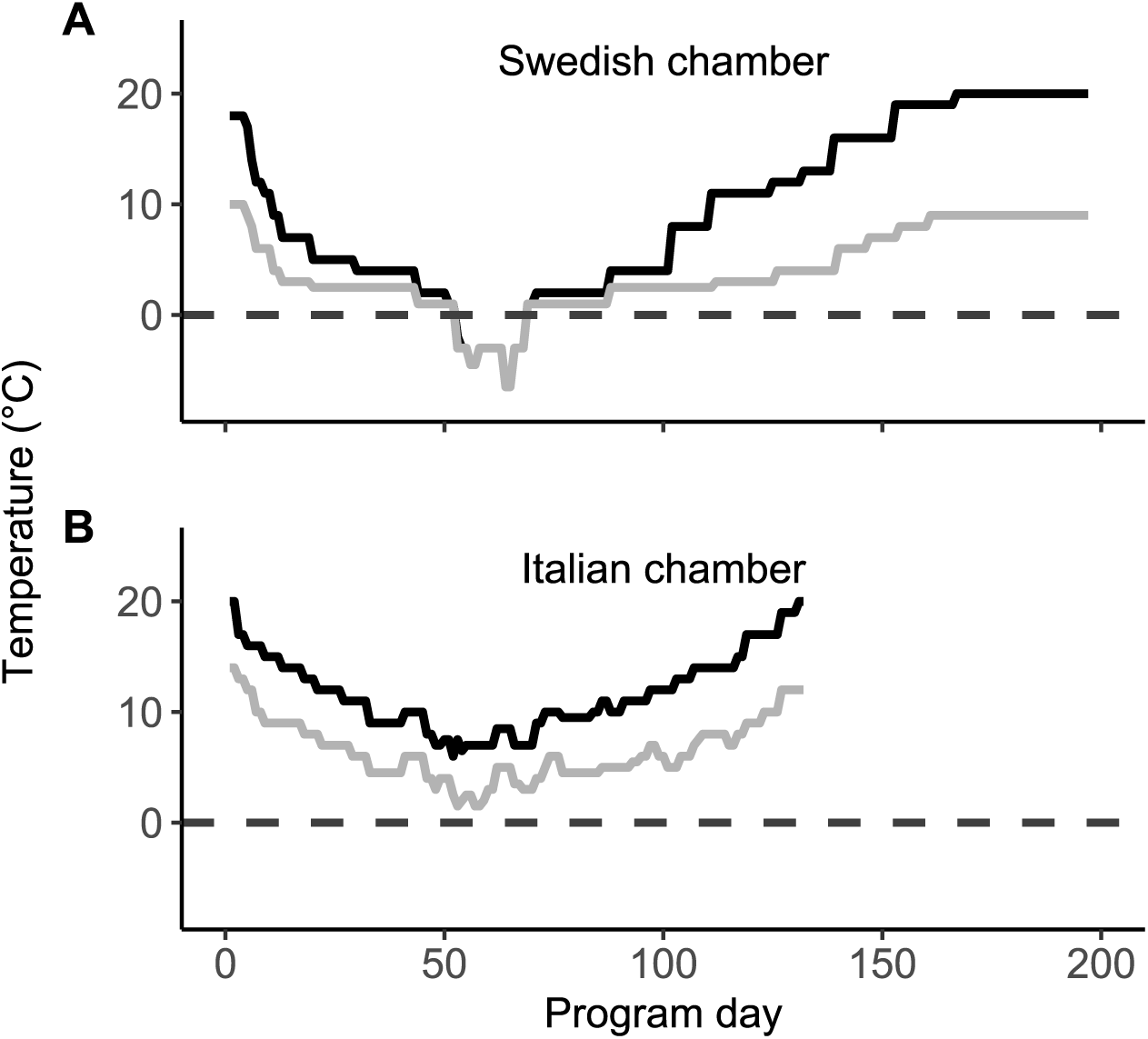
Temperature profiles for the growth chamber programs to mimic important aspects of the parental environments (Fig. S1). The programs were used for the Swedish growth chamber (A) and the Italian growth chamber (B) assays, respectively. Black lines indicate daily high temperatures and grey lines are daily low temperatures. Program day of the Italian chamber is shorter than the Swedish chamber because of the shorter life cycle at the native field site in Italy (see Figure S2).

**Figure S4.**
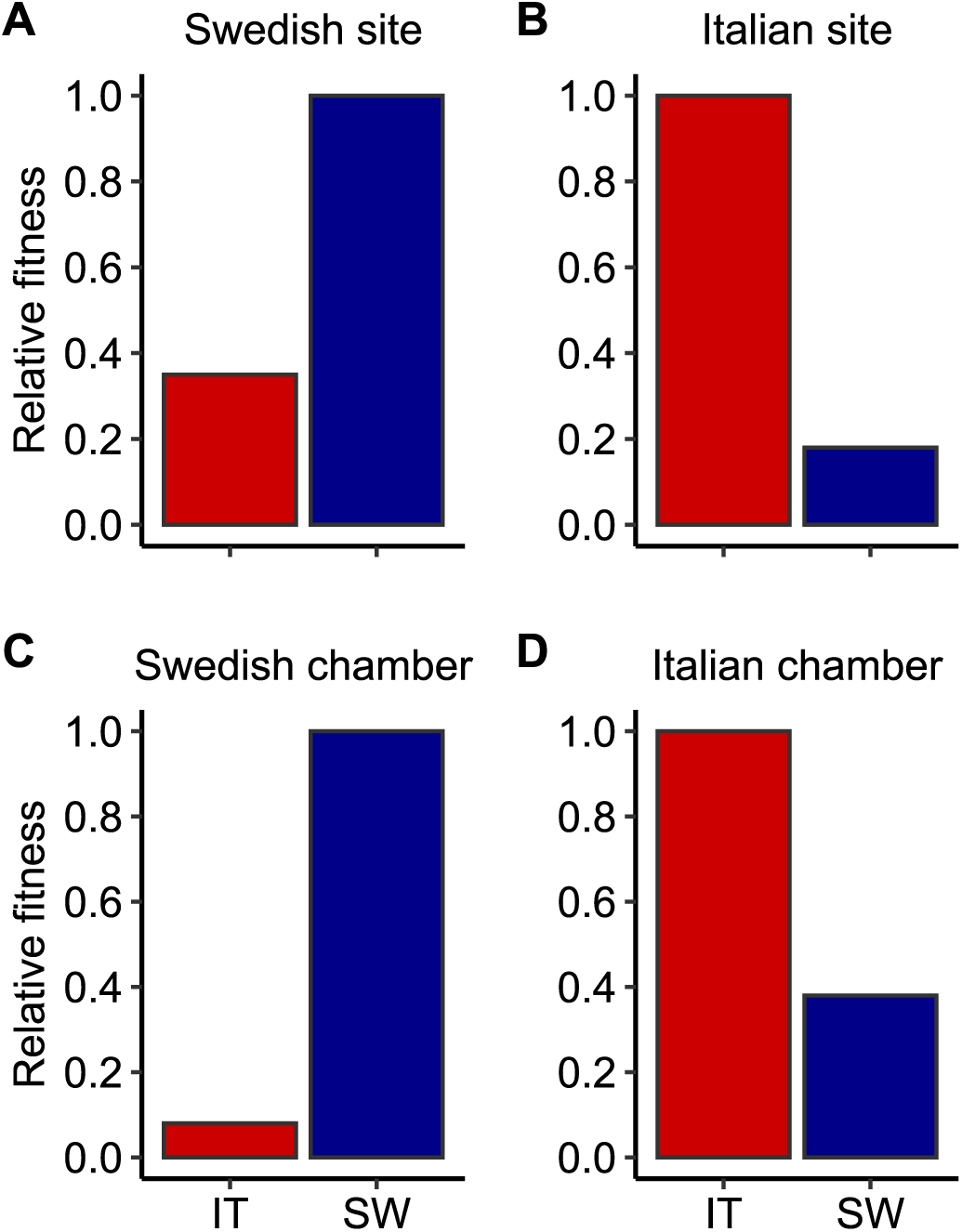
Comparison of relative fitness of the Italian (IT) and Swedish (SW) ecotypes between the field (A & B) and growth chamber (C & D) experiments.

**Figure S5.**
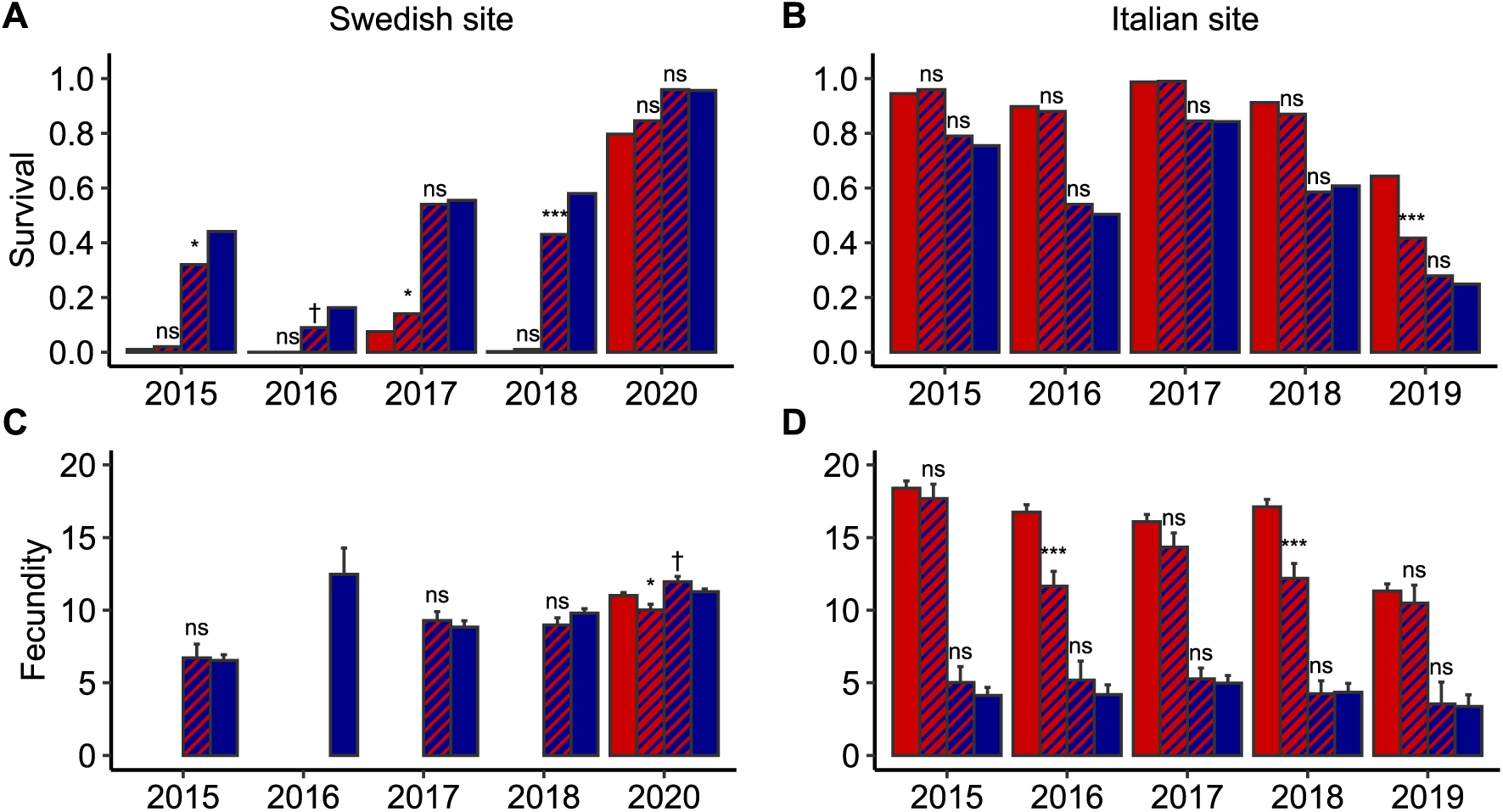
Fitness components of the SW and IT ecotypes and NILs separately for each of the five individual one-year experiments at the Swedish and Italian field sites. Mean proportion survival at the Swedish (A) and Italian (B) field sites. LSM fecundity at the same sites (C and D). Error bars for fecundity are 1 SE, no bars are given for survival because it was analyzed with a binomial error distribution. At the Swedish site, lines with 30 or fewer surviving individuals were omitted from analyses of fecundity, and these are represented by missing bars in C. Asterisks represent statistically significant contrasts between a NIL and its ecotypic genetic background (***, *P* < 0.001; **, *P* < 0.01; *, *P* < 0.05; ^†^, 0.05 < *P* < 0.1; ns, not significant).

**Figure S6.**
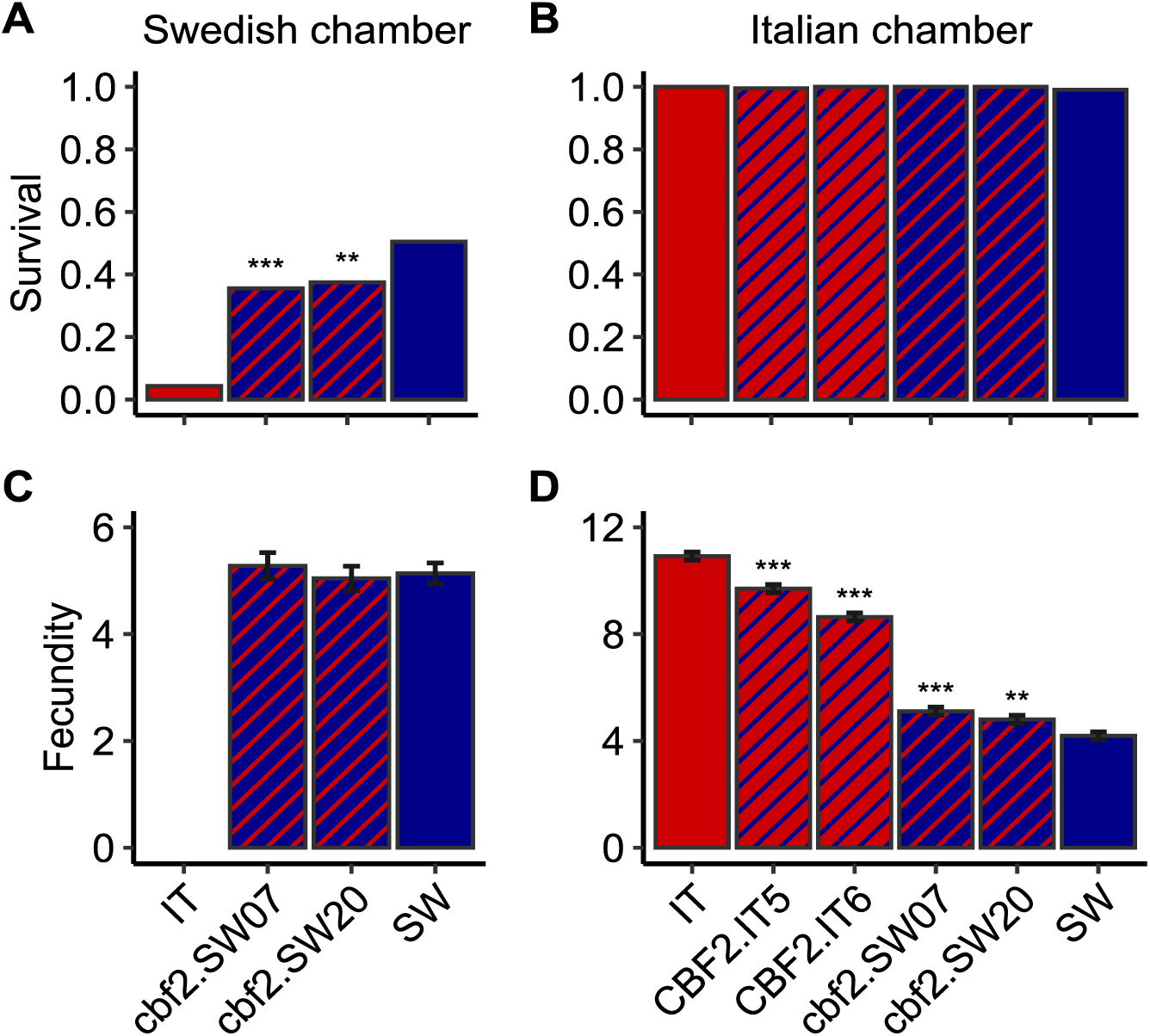
Fitness components of the SW and IT ecotypes and engineered lines in the Swedish and Italian chambers. Proportion survival (averaged over flats) in the Swedish (A) and Italian (B) chambers. LSM fecundity in the same chambers (C and D). Error bars for fecundity are 1 SE, no bars are given for survival because it was analyzed with a binomial error distribution. The Italian ecotype had fewer than 30 surviving individuals and it was excluded from fecundity analyses (missing bar in C). Asterisks represent statistically significant contrasts between an engineered line and its genetic background (***, *P* < 0.001; **, *P* < 0.01; *, *P* < 0.05; ^†^, 0.05 < *P* < 0.1; ns, not significant). No contrasts were performed for C because there was no significant main effect of genotype.

**Table S1.** ANOVA results for estimates of fitness and fitness components at the two field sites over 5 individual year experiments. For each dependent variable, the effects of genotype (2 ecotypes and 2 NILs), year, and the interaction between genotype and year were tested. Linear contrasts are also given between the IT and SW ecotypes over years and for each individual year. Because of near complete mortality in Sweden in 2019, results are presented for this year in Italy only. Data from an additional year (2020) were collected in Sweden to provide 5 years of data at each site. For fitness and fecundity, numerator degrees of freedom, F ratios, and P values are given as well as the t statistic and P values for the contrasts. For survival (binomially distributed), DF and *X*^2^ test statistics are given. The genotype by year interaction could not be included in the model for fecundity at the Swedish site because of high mortality of some lines in some years. In this case, contrasts between IT and SW are given for 2020 from a model run separately for this year.

|  | Swedish field site |  |  |  | Italian field site |  |  |
| --- | --- | --- | --- | --- | --- | --- | --- |
| **Fitness** | Effect | Num. DF | F | P | Num. DF | F | P |
|  | Genotype | 3 | 219.7 | **<0.001** | 3 | 736.8 | **<0.001** |
|  | Year | 4 | 538.8 | **<0.001** | 4 | 81.6 | **<0.001** |
|  | Genotype*Year | 12 | 11.2 | **<0.001** | 12 | 14.8 | **<0.001** |
|  | <u>IT vs. SW contrasts</u> |  | t | P |  | t | P |
|  | Overall |  | 23.7 | **<0.001** |  | 43.6 | **<0.001** |
|  | 2015 |  | 8.5 | **<0.001** |  | 23.0 | **<0.001** |
|  | 2016 |  | 6.1 | **<0.001** |  | 21.1 | **<0.001** |
|  | 2017 |  | 13.6 | **<0.001** |  | 18.9 | **<0.001** |
|  | 2018 |  | 16.9 | **<0.001** |  | 21.0 | **<0.001** |
|  | 2019 |  | - | - |  | 12.5 | **<0.001** |
|  | 2020 |  | 7.4 | **<0.001** |  | - | - |
| **Survival** | Effect | DF | $\chi^2$ | P | DF | $\chi^2$ | P |
|  | Genotype | 3 | 1004.4 | **<0.001** | 3 | 554.3 | **<0.001** |
|  | Year | 4 | 1957.3 | **<0.001** | 4 | 761.7 | **<0.001** |
|  | Genotype*Year | 12 | 99.7 | **<0.001** | 12 | 28.1 | **0.005** |
| | <u>IT vs. SW contrasts</u> | | $\chi^2$ | P | | $\chi^2$ | P |
|  | Overall |  | 126.8 | **<0.001** |  | 461.7 | **<0.001** |
|  | 2015 |  | 76.3 | **<0.001** |  | 47.7 | **<0.001** |
|  | 2016 |  | 12.6 | **<0.001** |  | 126.2 | **<0.001** |
|  | 2017 |  | 161.6 | **<0.001** |  | 33.3 | **<0.001** |
|  | 2018 |  | 51.2 | **<0.001** |  | 86.7 | **<0.001** |
|  | 2019 |  | - | - |  | 188.5 | **<0.001** |
|  | 2020 |  | 75.4 | **<0.001** |  | - | - |
| **Fecundity** | Effect | Num. DF | F | P | Num. DF | F | P |
|  | Genotype | 3 | 3.3 | **0.021** | 3 | 410.4 | **<0.001** |
|  | Year | 3 | 59.5 | **<0.001** | 4 | 9.8 | **<0.001** |
|  | Genotype*Year | - | - | - | 12 | 4.3 | **<0.001** |
|  | <u>IT vs. SW contrasts</u> |  | t | P |  | t | P |
|  | Overall |  | - | - |  | 32.3 | **<0.001** |
|  | 2015 |  | - | - |  | 19.2 | **<0.001** |
|  | 2016 |  | - | - |  | 15.0 | **<0.001** |
|  | 2017 |  | - | - |  | 15.6 | **<0.001** |
|  | 2018 |  | - | - |  | 60.0 | **<0.001** |
|  | 2019 |  | - | - |  | 8.4 | **<0.001** |
|  | 2020 |  | 0.9 | 0.349 |  | - | - |

**Table S2.** Effect sizes expressed as percent change relative to the background ecotype for the NILs (field experiments) and engineered lines (growth chamber experiments) for estimates of cumulative fitness and fitness components. Individual year estimates in Sweden were not calculated (nd) when one or more of the lines being compared had a survival probability of 2% or lower (fitness and survival) or had 30 or fewer surviving plants (fecundity). Effect sizes associated with contrast p values of < 0.1 are indicated in bold (see figures for details).

|  | Year | Line | Background | Fitness | Survival | Fecundity |
| --- | --- | --- | --- | --- | --- | --- |
| *Field sites* |  |  |  |  |  |  |
| *Sweden* |  |  |  |  |  |  |
|  | Overall | C_19 | IT | 4 | **15** | **-11** |
|  |  | R_38 | SW | **-12** | **-13** | 3 |
|  | 2015 | C_19 | IT | nd | nd | nd |
|  |  | R_38 | SW | -25 | **-28** | 3 |
|  | 2016 | C_19 | IT | nd | nd | nd |
|  |  | R_38 | SW | **-71** | **-45** | nd |
|  | 2017 | C_19 | IT | 147 | **87** | nd |
|  |  | R_38 | SW | 2 | -3 | 5 |
|  | 2018 | C_19 | IT | nd | nd | nd |
|  |  | R_38 | SW | **-32** | **-26** | -8 |
|  | 2020 | C_19 | IT | -3 | 6 | **9** |
|  |  | R_38 | SW | 6 | 0 | **6** |
| *Italy* |  |  |  |  |  |  |
|  | Overall | C_19 | IT | **-21** | -6 | **-17** |
|  |  | R_38 | SW | 14 | 3 | 11 |
|  | 2015 | C_19 | IT | -2 | 2 | -4 |
|  |  | R_38 | SW | 27 | 5 | 22 |
|  | 2016 | C_19 | IT | **-32** | -2 | **-31** |
|  |  | R_38 | SW | 33 | 7 | 24 |
|  | 2017 | C_19 | IT | **-11** | 0 | -11 |
|  |  | R_38 | SW | 6 | 0 | 6 |
|  | 2018 | C_19 | IT | **-32** | -5 | **-29** |
|  |  | R_38 | SW | -6 | -4 | -2 |
|  | 2019 | C_19 | IT | **-40** | **-35** | -7 |
|  |  | R_38 | SW | 18 | 12 | 5 |
| *Growth chambers* |  |  |  |  |  |  |
| *Sweden* |  |  |  |  |  |  |
|  |  | cbf2.SW07 | SW | **-23** | **-30** | 3 |
|  |  | cbf2.SW20 | SW | **-22** | **-26** | -2 |
| *Italy* |  |  |  |  |  |  |
|  |  | CBF2.IT5 | IT | **-12** | -1 | **-11** |
|  |  | CBF2.IT6 | IT | **-21** | 0 | **-21** |
|  |  | cbf2.SW07 | SW | **23** | 1 | **22** |
|  |  | cbf2.SW20 | SW | **16** | 1 | **15** |

**Table S3.**
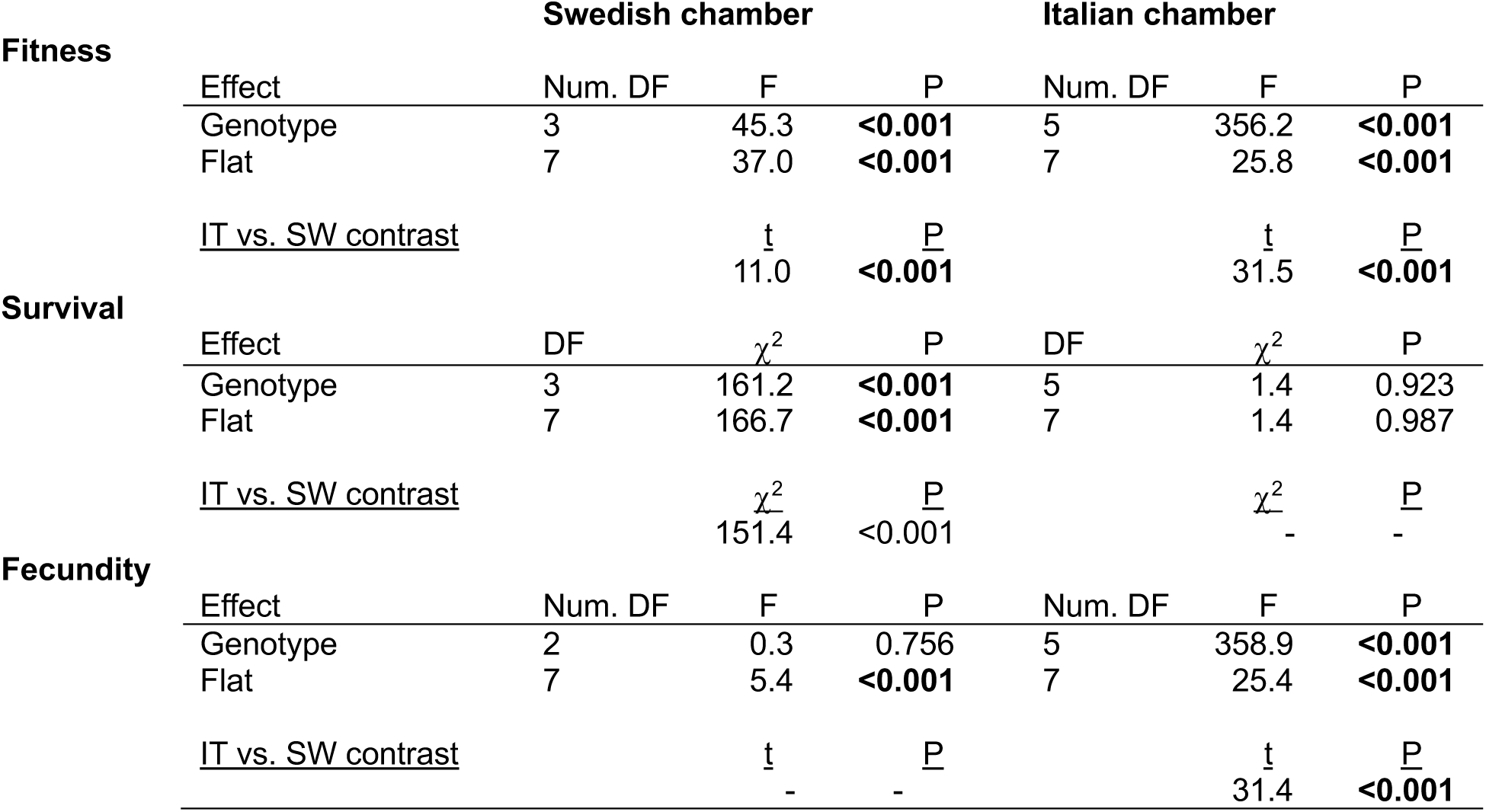
ANOVA results for estimates of fitness and fitness components in the growth chamber experiments simulating the Swedish and Italian environments. For each dependent variable the effects of genotype (2 ecotypes and 2 CRISPR lines in the Swedish growth chamber, and an additional 2 transgenic lines in the Italian growth chamber) were tested. We also include a term for tray to account for spatial variation within the chamber. Linear contrasts are given between the IT and SW ecotypes. For fitness and fecundity, numerator degrees of freedom, F ratios, and P values are given as well as the t statistic and P values for the contrasts. For survival (binomially distributed), DF and *X*^2^ test statistics are given. In cases where there was not a significant effect of genotype (survival in the Italian chamber, fecundity in the Swedish chamber), no contrasts were performed.

## Notes

### Competing Interest Statement

The authors have declared no competing interest.

### Summary of Updates

Revised text in results and discussion, added a new supplemental table (not Table S2).

## References

1. Kawecki TJ & Ebert D (2004) Conceptual issues in local adaptation. Ecology Letters 7:1225–1241.

2. Hereford J (2009) A quantitative survey of local adaptation and fitness trade-offs. The American Naturalist 173:579–588.

3. MacArthur RH (1972) Geographical Ecology: Patterns in the Distribution of Species. (Princeton University Press, Princeton, New Jersey).

4. Futuyma DJ & Moreno G (1988) The evolution of ecological specialization. Annual Review of Ecology and Systematics 19:207–233.

5. Agrawal AA (2020) A scale-dependent framework for trade-offs, syndromes, and specialization in organismal biology. Ecology 101:e02924.

6. Leimu R & Fischer M (2008) A meta-analysis of local adaptation in plants. Public Library of Science ONE 3:10.1371/journal.pone.0004010.t0004001.

7. Johnson LC, et al. (2022) Reciprocal transplant gardens as gold standard to detect local adaptation in grassland species: New opportunities moving into the 21st century. Journal of Ecology 110:1054–1071.

8. VanWallendael A, Lowry DB, & Hamilton JA (2022) One hundred years into the study of ecotypes, new advances are being made through large-scale field experiments in perennial plant systems. Current Opinion in Plant Biology 66:102152.

9. Savolainen O, Lascoux M, & Merilä J (2013) Ecological genomics of local adaptation. Nature Reviews Genetics 14(11):807–820.

10. Tiffin P & Ross-Ibarra J (2014) Advances and limits of using population genetics to understand local adaptation. Trends in Ecology & Evolution 29(12):673–680.

11. VanWallendael A, et al. (2019) A Molecular View of Plant Local Adaptation: Incorporating Stress-Response Networks. Annual Review of Plant Biology 70:14.11–14.25.

12. Wadgymar SM, DeMarche ML, Josephs EB, Sheth SN, & Anderson JT (2022) Local adaptation: Causal agents of selection and adaptive trait divergence. Annual Review of Ecology, Evolution, and Systematics 53:87–111.

13. Stinchcombe JR & Hoekstra HE (2008) Combining population genomics and quantitative genetics: finding the genes underlying ecologically important traits. Heredity 100:158–170.

14. Barrett RDH & Hoekstra HE (2011) Molecular spandrels: tests of adaptation at the genetic level. Nature Reviews Genetics 12:767–780.

15. Bomblies K & Peichel CL (2022) Genetics of adaptation. Proceedings of the National Academy of Sciences U S A 119(30):e2122152119.

16. Rockman MV (2012) The QTN program and the alleles that matter for evolution: all that’s gold does not glitter. Evolution 66:1–17.

17. Rausher MD & Delph LF (2015) Commentary: When does understanding phenotypic evolution require identification of the underlying genes? Evolution 69:1655–1664.

18. Dittmar EL, Oakley CG, Conner JK, Gould BA, & Schemske DW (2016) Factors influencing the effect size distribution of adaptive substitutions. Proceedings of the Royal Society B-Biological Sciences 283:20153065.

19. Schluter D, et al. (2021) Fitness maps to a large-effect locus in introduced stickleback populations. Proceedings of the National Academy of Sciences 118:e1914889118.

20. Anderson JT, Willis JH, & Mitchell-Olds T (2011) Evolutionary genetics of plant adaptation. Trends in Genetics 27:258–266.

21. Wadgymar SM, et al. (2017) Identifying targets and agents of selection: innovative methods to evaluate the processes that contribute to local adaptation. Methods in Ecology and Evolution 8:738–749.

22. Lowry DB, Hall MC, Salt DE, & Willis JH (2009) Genetic and physiological basis of adaptive salt tolerance divergence between coastal and inland *Mimulus guttatus*. New Phytologist 183(3):776–788.

23. Hall MC, Lowry DB, & Willis JH (2010) Is local adaptation in *Mimulus guttatus* caused by trade-offs at individual loci? Molecular Ecology 19(13):2739–2753.

24. Anderson JT, Lee C-R, Rushworth CA, Colautti RI, & Mitchell-Olds T (2013) Genetic trade-offs and conditional neutrality contribute to local adaptation. Molecular Ecology 22(3):699–708.

25. Leinonen PH, Remmington DL, Leppälä J, & Savolainen O (2013) Genetic basis of local adaptation and flowering time variation in *Arabidopsis lyrata*. Molecular Ecology 22(3):709–723.

26. Ågren J, Oakley CG, McKay JK, Lovell JT, & Schemske DW (2013) Genetic mapping of adaptation reveals fitness tradeoffs in *Arabidopsis thaliana*. Proceedings of the National Academy of Sciences of the United States of America 110:21077–21082.

27. Postma FM & Ågren J (2016) Early life stages contribute strongly to local adaptation in *Arabidopsis thaliana*. Proceedings of the National Academy of Sciences of the United States of America 113:7590–7595.

28. Wright SJ, Goad DM, Gross BL, Muñoz PR, & Olsen KM (2021) Genetic trade-offs underlie divergent life history strategies for local adaptation in white clover. Molecular Ecology:10.1111/mec.16180.

29. Oakley CG, Schemske DW, McKay JK, & Ågren J (2023) Ecological genetics of local adaptation in Arabidopsis: An 8-year field experiment. Molecular Ecology 32:4570–4583.

30. Mackay TFC, Stone EA, & Ayroles JF (2009) The genetics of quantitative traits: challenges and prospects. Nature Reviews Genetics 10:565–577.

31. Des Marais DL, Hernandez KM, & Juenger TE (2013) Genotype-by-environment interaction and plasticity: exploring genomic responses of plants to the abiotic environment. Annual Review of Ecology, Evolution, and Systematics 44:5–29.

32. Alonso-Blanco C & Méndez-Vigo B (2014) Genetic architecture of naturally occuring quantitative traits in plants: an updated synthesis. Current Opinion in Plant Biology 18:37–43.

33. Des Marais DL, et al. (2014) Variation in *MPK12* affects water use efficiency in *Arabidopsis* and reveals a pleiotropic link between guard cell size and ABA response. Proceedings of the National Academy of Sciences of the United States of America 111(7):2836–2841.

34. Josephs EB (2018) Determining the evolutionary forces shaping G x E. New Phytologist 219:31–36.

35. Storey KB & Storey JM (1996) Natural freezing survival in animals. Annual Review of Ecology, Evolution, and Systematics 27:365–386.

36. Addo-Bediako A, Chown SL, & Gaston KJ (2000) Thermal tolerance, climatic variability and latitude. Proceedings of the Royal Society B-Biological Sciences 267:739–745.

37. Preston JC & Sandve SR (2013) Adaptation to seasonality and the winter freeze. Frontiers in Plant Science 4:10.3389/fpls.2013.00167.

38. Chang CY, Bräutigam K, Hüner NPA, & Ensminger I (2021) Champions of winter survival: cold acclimation and molecular regulation of cold hardiness in evergreen conifers. New Phytologist 229:675–691.

39. Levitt J (1980) Chilling, freezing, and high temperature stresses (Academic press, New York).

40. Colinet H, Larvor V, Laparie M, & Renault D (2012) Exploring the plastic response to cold acclimation through metabolomics. Functional Ecology 26:711–722.

41. Long Y, et al. (2013) Transcriptomic characterization of cold acclimation in larval zebrafish. BMC Genomics 14:612.

42. Kristensen TN, Kjeldal H, Schou MF, & Nielsen JL (2016) Proteomic data reveal a physiological basis for costs and benefits associated with thermal acclimation. Journal of Experimental Biology 219:969–976.

43. MacMillan HA, et al. (2016) Cold acclimation wholly reorganizes the *Drosophila melanogaster* transcriptome and metabolome. Scientific Reports 6:28999.

44. Noer NK, et al. (2022) Rapid adjustments in thermal tolerance and the metabolome to daily environmental changes – A field study on the arctic seed bug *Nysius groenlandicus*. Frontiers in Physiology 13:818485.

45. Thomashow MF (2010) Molecular basis of plant cold acclimation: insights gained from studying the CBF cold response pathway. Plant Physiology 154:571–577.

46. Loehle C (1998) Height growth tradeoffs determine northern and southern range limits for trees. Journal of Biogeography 25:735–742.

47. Auld JR, Agrawal AA, & Reylea RA (2010) Re-evauluating the costs and limits of adaptive pehotypic plasticity. Proceedings of the Royal Society B-Biological Sciences 277:503–511.

48. Kristensen TN, et al. (2008) Costs and benefits of cold acclimation in field-released Drosophila. Proceedings of the National Academy of Sciences of the United States of America 105:216–221.

49. Schou MF, Loeschcke V, & Kristensen TN (2015) Strong costs and benefits of winter acclimatization in *Drosophila melanogaster*. PLos ONE 10:e0130307.

50. Everman ER, Delzeit JL, Hunter FK, Gleason JM, & Morgan TJ (2018) Costs of cold acclimation on survival and reproductive behavior in *Drosophila melanogaster*. PLos ONE 13:e0197822.

51. Taji T, et al. (2002) Important roles of drought– and cold-inducible genes for galactinol synthase in stress tolerance in *Arabidopsis thaliana*. The Plant Journal 29:417–426.

52. Kaplan F, et al. (2007) Transcript and metabolite profiling during cold acclimation of *Arabidopsis* reveals an intricate relationship of cold-regulated gene expression with modifications in metabolite content. The Plant Journal 50:967–981.

53. Weiszmann J, et al. (2023) Metabolome plasticity in 241 Arabisopsis thaliana accessions reveals evolutionary cold adaptation process. Plant Physiology: 10.1093/plphys/kiad1298.

54. Barrero-Gil J & Salinas J (2018) Gene regulatory networks mediating cold acclimation: the CBF pathway. Survival Strageies in Extreme Cold and Desiccation, eds Iwaya-Inoue M, Sakurai M, & Uemura M (Springer Singapore, Singapore).

55. Park S, et al. (2015) Regulation of the *Arabidopsis* CBF regulon by a complex low-temperature regulatory network. The Plant Journal 82:193–207.

56. Alonso-Blanco C, et al. (2005) Genetic and molecular analyses of natural variation indicate *CBF2* as a candidate gene for underlying a freezing tolerance quantitative trait locus in *Arabidopsis*. Plant Physiology 139:1304–1312.

57. Oakley CG, Ågren J, Atchison RA, & Schemske DW (2014) QTL mapping of freezing tolerance: links to fitness and adaptive trade-offs. Molecular Ecology 23:4304–4315.

58. Gehan MA, et al. (2015) Natural variation in the C-repeat binding factor cold response pathway correlates with local adaptation of *Arabidopsis* ecotypes. The Plant Journal 84:682–693.

59. Sanderson BJ, et al. (2020) Genetic and physiological mechanisms of freezing tolerance in locally adapted populations of a winter annual. American Journal of Botany 107(2):250–261.

60. Savitch LV, et al. (2005) The effect of overexpression of two *Brassica CBF/DREB1*-like transcription factors on photosynthetic capacity and freezing tolerance in Brassica napus. Plant Cell Physiolology 46:1525–1539.

61. Vágújfalvi A, Galiba G, Cattivelli L, & Dubcovsky J (2003) The cold-regulated transcriptional activator *Cbf3* is linked to the frost-tolerance locus *Fr-A2* on wheat chromosome 5A. Molecular Genetics and Genomics 269:60–67.

62. Benedict C, et al. (2006) The *CBF1*-dependent low temperature signalling pathway, regulon and increase in freeze tolerance are conserved in *Populus* spp. Plant, Cell & Environment 29:1259–1272.

63. Savage JA & Cavender-Bares J (2013) Phenological cues drive an apparent trade-off between freezing tolerance and growth in the family Salicaceae. Ecology 94:1708–1717.

64. Jackson MW, Stinchcombe JR, Korves TM, & Schmitt J (2004) Costs and benefits of cold tolerance in transgenic *Arabidopsis thaliana*. Molecular Ecology 13:3609–3615.

65. Zhen Y, Dhakal P, & Ungerer MC (2011) Fitness benefits and costs of cold acclimation in Arabidopsis thaliana. American Naturalist 178(1):44–52.

66. Zhen Y & Ungerer MC (2008) Relaxed selection on the *CBF*/*DREB1* regulatory genes and reduced freezing tolerance in the southern range of *Arabidopsis thaliana*. Molecular Biology and Evolution 25(12):2547–2555.

67. Zuther E, Schulz E, Childs LH, & Hincha DK (2012) Clinal variation in the non-acclimated and cold-acclimated freezing tolerance of *Arabidopsis thaliana* accessions. Plant, Cell and Environment 35(10):1860–1878.

68. Monroe JG, et al. (2016) Adaptation to warmer climates by parallel functional evolution of CBF genes in *Arabidopsis thaliana*. Molecular Ecology 25:3632–3644.

69. Boinot M, Karakas E, Koehl K, Pagter M, & Zuther E (2022) Cold stress and freezing tolerance negatively affect the fitness of *Arabidopsis thaliana* accessions under field and controlled conditions. Planta 255:39.

70. Ågren J & Schemske DW (2012) Reciprocal transplants demonstrate strong adaptive differentiation of the model organism *Arabidopsis thaliana* in its native range. New Phytologist 194:1112–1122.

71. Zacchello G, Vinyeta M, & Ågren J (2020) Strong stabilizing selection on timing of germination in a Mediterranean population of *Arabidopsis thaliana*. American Journal of Botany.

72. Beavis WD (1994) The power and deceit of QTL experiments: lessons from comparative QTL studies. Proceedings of the Corn and Sorghum Industry Research Conference, American Seed Trade Association, Washington DC:250-266.

73. Martínez-Berdeja A, et al. (2020) Functional variants of *DOG1* control seed chilling responses and variation in seasonal life-history strategies in *Arabidopsis thaliana*. Proceedings of the National Academy of Sciences 117:2526–2534.

74. Wang J, et al. (2018) A major locus controls local adaptation and adaptive life history variation in a perennial plant. Genome Biology 19:72.

75. Taylor MA, et al. (2019) Large-effect flowering time mutations reveal conditionally adaptive paths through fitness landscapes in *Arabidopsis thaliana*. Proceedings of the National Academy of Sciences 116:17890–17899.

76. Lopez-Arboleda WA, Reinert S, Nordborg M, & Korte A (2021) Global genetic heterogeneity in adaptive traits. Molecular Biology and Evolution 38:4822–4831.

77. Fournier-Level A, et al. (2011) A map of local adaptation in *Arabidopsis thaliana*. Science 334(6052):86–89.

78. Hancock AM, et al. (2011) Adaptation to climate across the *Arabidopsis thaliana* genome. Science 334(6052):83–86.

79. Exposito-Alonso M, et al. (2019) Natural selection on the *Arabidopsis thaliana* genome in present and future climates. Nature 573:126–129.

80. Prasad KV, et al. (2012) A gain-of-function polymorphism controlling complex traits and fitness in nature. Science 337:1081–1084.

81. Barrett RDH, et al. (2019) Linking a mutation to survival in wild mice. Science 363:499–504.

82. Kerwin R, et al. (2015) Natural genetic variation in *Arabidopsis thaliana* defense metabolism genes modulates field fitness. Elife 4:10.7554/eLife.05604.

83. Zhen Y & Ungerer MC (2008) Clinal variation in freezing tolerance among natural accessions of *Arabidopsis thaliana*. New Phytologist 177(2):419–427.

84. Koornneef M, Alonso-Blanco C, & Vreugdenhil D (2004) Naturally occurring genetic variation in *Arabidopsis thaliana*. Annual Review of Plant Biology 55:141–172.

85. Durvasula A, et al. (2017) African genomes illuminate the early history and transition to selfing in *Arabidopsis thaliana*. *Proceedings of the National Academy of Sciences*, USA 114:5213–5218.

86. Price N, et al. (2018) Combining population genomics and fitness QTLs to identify the genetics of local adaptation in *Arabidopsis thaliana*. Proceedings of the National Academy of Sciences 115(19):5028–5033.

87. Ellis TJ, Postma FM, Oakley CG, & Ågren J (2021) Life-history trade-offs and the genetic basis of fitness in *Arabidopsis thaliana*. Molecular Ecology 30:2846–2858.

88. Ågren J, Oakley CG, Lundemo S, & Schemske DW (2017) Adaptive divergence in flowering time among natural populations of *Arabidopsis thaliana*: estimates of selection and QTL mapping. Evolution 71:550–564.

89. Oakley CG, et al. (2018) Genetic basis of photosynthetic responses to cold in two locally adapted populations of *Arabidopsis thaliana*. Journal of Experimental Botany 69:699–709.

90. Cohu CM, Muller O, Stewart JJ, Demmig-Adams B, & Adams WWI (2013) Association between minor loading vein architecture and light– and CO2-saturated rates of photosynthetic oxygen evolution among *Arabidopsis thaliana* ecotypes from different latitudes. Frontiers in Plant Science 4:240.

91. Stewart JJ, et al. (2015) Differences in light-harvesting, acclimation to growth-light environment, and leaf structural development between Swedish and Italian ecotypes of *Arabidopsis thaliana*. Planta 242(6):1277–1290.

## SI References

1. Ågren J & Schemske DW (2012) Reciprocal transplants demonstrate strong adaptive differentiation of the model organism *Arabidopsis thaliana* in its native range. New Phytologist 194:1112–1122.

2. Oakley CG, Ågren J, Atchison RA, & Schemske DW (2014) QTL mapping of freezing tolerance: links to fitness and adaptive trade-offs. Molecular Ecology 23:4304–4315.

3. Dittmar EL, Oakley CG, Ågren J, & Schemske DW (2014) Flowering time QTL in natural populations of *Arabidopsis thaliana* and implications for their adaptive value. Molecular Ecology 23:4291–4303.

4. Gehan MA, Greenham K, Mockler TC, & McClung CR (2015) Transcriptional networks — crops, clocks, and abiotic stress. Current Opinion in Plant Biology 24:39–46.

5. Dong MA, Farre EM, & Thomashow MF (2011) CIRCADIAN CLOCK-ASSOCIATED 1 and LATE ELONGATED HYPOCOTYL regulate expression of the C-REPEAT BINDING FACTOR (CBF) pathway in *Arabidopsis*. PNAS 108:7241–7246.

6. Zacchello G, Vinyeta M, & Ågren J (2020) Strong stabilizing selection on timing of germination in a Mediterranean population of *Arabidopsis thaliana*. American Journal of Botany.

7. Oakley CG, Schemske DW, McKay JK, & Ågren J (2023) Ecological genetics of local adaptation in Arabidopsis: An 8-year field experiment. Molecular Ecology.

